## Supplementary File 1 for "APOGEE 2: multi-layer machine-learning model for the interpretable prediction of mitochondrial missense variants"

### MD PROTOCOL

**# Minimization and equilibration protocol generated by CHARMM-GUI (http://www.charmm-gui.org) v3.5**
**# GaMD production protocol implemented following [https://miaolab.ku.edu/GaMD/manual.html](https://miaolab.ku.edu/GaMD/manual.html) guidelines**
**# All commands/inputs were implemented for AMBER20, so a lower version of AMBER can cause some errors.**

```
set amber = pmemd

set init = step5_input
set mini_prefix = step6.0_minimization
set equi_prefix = step6.%d_equilibration
set prod_prefix = step7_production
set prod_step   = step7
```

**# Minimization**

```
if (-e dihe.restraint) sed -e "s/FC/250.0/g" dihe.restraint > ${mini_prefix}.rest
pmemd -O -i ${mini_prefix}.mdin -p ${init}.parm7 -c ${init}.rst7 -o ${mini_prefix}.mdout
-r ${mini_prefix}.rst7 -inf ${mini_prefix}.mdinfo -ref ${init}.rst7
```

**# Equilibration**

```
set cnt = 1
set cntmax = 6
set fc = {'250.0','100.0','50.0','50.0','25.0'}

while ( ${cnt} <= ${cntmax} )
       @ pcnt = ${cnt} - 1
       set istep = `printf ${equi_prefix} ${cnt}`
       set pstep = `printf ${equi_prefix} ${pcnt}`
       if ( ${cnt} == 1 ) set pstep = ${mini_prefix}

       if (-e dihe.restraint && ${cnt} < ${cntmax}) sed -e "s/FC/${fc[${cnt}]}/g"
dihe.restraint > ${istep}.rest
       ${amber} -O -i ${istep}.mdin -p ${init}.parm7 -c ${pstep}.rst7 -o ${istep}.mdout -
r ${istep}.rst7 -inf ${istep}.mdinfo -ref ${init}.rst7 -x ${istep}.nc
       @ cnt += 1
end
```

**# Gaussian accelerated Molecular Dynamics production**

**#Run initial conventional molecular dynamics and, after ntcmd+ntebsteps (defined in prep_gamd.in), run GaMD equilibration**

```
pmemd.cuda -O -i prep_gamd.in -o 1_gamd_Prod.out -p step5_input.parm7 -c
step6.6_equilibration.rst7 -inf 1_gamd_Prod.info -x 1_gamd_Prod.mdcrd -r 1_gamd_Prod.rst
-gamd gamd_prod_prep.log
```

**#start GaMD production simulation**

```
pmemd.cuda -O -i 1_gamd.in -o 2_gamd_Prod.out -p step5_input.parm7 -c 1_gamd_Prod.rst -
inf 2_gamd_Prod.info -x 2_gamd_Prod.mdcrd -r 2_gamd_Prod.rst -gamd gamd_prod_1.log
```

**#repeat running jobs using the same input file (${cnt}_gamd.in) until end of the GaMD production simulation**

```
set cnt    = 3
set cntmax = 10
while ( ${cnt} <= ${cntmax} )
       pmemd.cuda -O -i  ${cnt}_gamd.in -o  ${cnt}_gamd_Prod.out -p step5_input.parm7 -
c  $((cnt-1))_gamd_Prod.rst -inf  ${cnt}_gamd_Prod.info -x  ${cnt}_gamd_Prod.mdcrd -
r  ${cnt}_gamd_Prod.rst -gamd gamd_prod_${cnt}.log
       @ cnt += 1
end
```

**#Template input file (prep_gamd.in, 1_gamd.in, 2-10_gamd.in) for GaMD production steps**

**prep_gamd.in**
```
&cntrl
imin = 0,
irest = 0,
ntx = 1,
ioutfm = 1,
nstlim = 50000000,
dt = 0.002,
ntt = 3,
gamma_ln = 5.0,
ig = -1,
tempi = 300.0,
temp0 = 300.0,
ntp = 0,
ntb = 1,
ntc = 2,
ntf = 2,
cut = 12,
ntwr = 500,
ntpr = 200,
ntwx = 5000,
ntwe = 200,
iwrap = 1,
ntr = 0,
ntwprt = 178474,

igamd = 3, iE = 1, irest_gamd = 0,
ntcmd = 10000000, nteb = 50000000, ntave =2000000,
ntcmdprep = 4000000, ntebprep = 4000000,
sigma0P = 12.0, sigma0D = 12.0,
&end
```

**1_gamd.in**
```
&cntrl
imin = 0,
irest = 0,
ntx = 1,
ioutfm = 1,
nstlim = 10000000,
dt = 0.002,
ntt = 3,
gamma_ln = 5.0,
ig = -1,
tempi = 300.0,
temp0 = 300.0,
ntp = 0,
ntb = 1,
ntc = 2,
ntf = 2,
cut = 12,
ntwr = 500,
ntpr = 200,
ntwx = 400,
ntwe = 200,
iwrap = 1,
ntr = 0,
ntwprt = 178474,

igamd = 3, iE = 1, irest_gamd = 1,
ntcmd = 0, nteb = 0, ntave = 2000000,
ntcmdprep = 0, ntebprep = 0,
sigma0P = 12.0, sigma0D = 12.0,
&end
```

**2-10_gamd.in**

```
&cntrl
imin = 0,
irest = 1,
ntx = 5,
ioutfm = 1,
nstlim = 10000000,
dt = 0.002,
ntt = 3,
gamma_ln = 5.0,
ig = -1,
tempi = 300.0,
temp0 = 300.0,
ntp = 0,
ntb = 1,
ntc = 2,
ntf = 2,
cut = 12,
ntwr = 500,
ntpr = 200,
ntwx = 400,
ntwe = 200,
iwrap = 1,
ntr = 0,
ntwprt = 178474,

igamd = 3, iE = 1, irest_gamd = 1,
ntcmd = 0, nteb = 0, ntave = 2000000,
ntcmdprep = 0, ntebprep = 0,
sigma0P = 12.0, sigma0D = 12.0,
&end
```
